# Incidental learning of predictive temporal context within cortical representations of visual shape

**DOI:** 10.1101/2023.05.19.541422

**Authors:** Ehsan Kakaei, Jochen Braun

**Affiliations:** European Structural and Investment Funds Graduate School on Analysis, Imaging, and Modelling of Neuronal and Inflammatory Processes, Otto-von-Guericke University, 39120 Magdeburg, Germany; Institute of Biology, Otto-von-Guericke University, 39120 Magdeburg, Germany; Center for Behavioral Brain Sciences, Otto-von-Guericke University, 39120 Magdeburg, Germany

## Abstract

Incidental learning of spatio-temporal regularities and consistencies – also termed ‘statistical learning’ – may be important for discovering the causal principles governing the world. We studied statistical learning of temporal structure at two temporal scales: the presentation of rotating visual objects (3 s) and predictive temporal context (30 s) in a sequence of such objects.

Observers viewed fifteen initially unfamiliar synthetic objects recurring many times each, intermixed with other similar objects that appeared only once, while whole-brain BOLD activity was recorded. Analyzing multivariate BOLD activity, we located view-independent responses to synthetic objects throughout the ventral occipitotemporal cortex, revealing a view-invariant representation of object shape.

Predictive temporal context modified representations of object shape at early and middle levels of the visual processing hierarchy. Approximately 20% of the sites representing shape also exhibited a significant representation of temporal context. A few higher-level sites represented temporal context, without also representing object shape. Additionally, objects marking transitions between communities were distinctly represented at many sites.

We conclude that, at the sensitivity afforded by fMRI, the cortical representations of invariant visual shape and of predictive temporal context are largely coextensive. Our results suggest that incidental learning of causal principles and structures operating at different timescales (“structural learning”) may involve visually responsive areas in the ventral occipitotemporal cortex.

## 1 Introduction

Even when sensory stimuli are experienced passively – without task or reward – they can modify the underlying neural pathways and alter subsequent sensory performance and behavior (*e.g.*, Conway and Christiansen 2005; Li and DiCarlo 2012; Lengyel et al. 2019). This incidental and automatic type of plasticity had been termed ‘statistical learning’ or ‘implicit learning’ (for reviews, see Perruchet and Pacton 2006; Aslin 2017; Schapiro and Turk-Browne 2015; Saffran and Kirkham 2018; Perruchet 2019; Fiser and Lengyel 2022). Some theories of cognitive development hypothesize that incidental learning during everyday experience captures the causal processes and relationships underlying sensory observations at a more abstract level (Kemp and Tenenbaum, 2008; Tenenbaum et al., 2011). If so, statistical learning might contribute to higher cognitive function by acquiring the quality of a “structural learning” that could underpin learning from examples, generalizing between domains, or gaining causal insight and understanding (Shafto et al., 2011; Lake et al., 2017).

A well-studied instance of incidental learning is the view-invariance of visual object recognition (for reviews, see Logothetis and Sheinberg 1996; DiCarlo et al. 2012; Gauthier and Tarr 2016). Humans and non-human primates typically recognize visual objects from different viewing directions and distances, presumably relying on characteristic features and/or their spatial relationships. This perceptual invariance can be modified rapidly by the experience of contiguous sequences of different views, demonstrating dependence on learning (Wallis and Bülthoff, 2001; Wallis et al., 2009; Tian and Grill-Spector, 2015). The neural representation of visual shape in the ventral occipitotemporal cortex is similarly view-invariant and equally subject to modification by the recent experience of (natural or unnatural) sequences of views (Li and DiCarlo, 2008, 2010, 2012; de Beeck and Baker, 2010; Van Meel and Op de Beeck, 2018; Van Meel and de Beeck, 2020; Jia et al., 2021).

Incidental learning is not limited to individual objects but extends also to spatiotemporal configurations of multiple objects. When human observers experience temporal sequences or spatial arrays of visual objects, task-irrelevant statistical regularities and contingencies are learned rapidly (within minutes), as can be revealed by subsequent behavioral tests (Fiser and Aslin, 2001, 2002, 2005; Turk-Browne et al., 2005, 2009; Kakaei et al., 2021; Śaringer et al., 2022). In non-human primates, the experience of task-irrelevant temporal dependencies modifies object-specific responses of neurons in visual areas of the ventral temporal cortex, but also in multimodal areas of the medial temporal lobe (Miyashita 1988; Sakai and Miyashita 1991 **AIT**; Erickson and Desimone 1999 **perirhinal**; Meyer et al. 2014 **IT**; Kaposvari et al. 2018 **IT**). In human observers, functional imaging evidence reveals that task-irrelevant temporal dependen-cies can modulate BOLD responses in visually selective areas of the ventral occipital cortex, as well as in multimodal areas such as the medial temporal lobe, hippocampus, and basal ganglia (Turk-Browne et al. 2009 **LOC, VOTC**; Turk-Browne et al. 2010 **anterior HC**; Gheysen et al. 2011 **caudate, HC**; Schapiro et al. 2012 **HC**; Hsieh et al. 2014 **HC**;Hindy et al. 2016 **HC**; Wang et al. 2017; Giorgio et al. 2018; Karlaftis et al. 2019 **frontal-cingulate-sensorymotor-basal ganglia vs. occipitotemporal-hippocampus-basal ganglia**). Statistical learning goes beyond first-order dependencies (between immediate temporal neighbors) and extends to higher-order dependencies (between more distant neighbors). For example, Schapiro and colleagues demonstrated statistical learning of clusters of dependencies (“temporal communities”) and observed BOLD correlates of this predictive temporal context in associative areas of the frontal and temporal lobes and in the hippocampus, but not in visually selective areas of the ventral occipitotemporal cortex (Schapiro et al. 2013 **inferior frontal gyrus, insula, anterior temporal lobe and superior temporal gyrus**;Schapiro et al. 2016 **HC**).

Here we investigate statistical learning by human observers with temporal sequences of visual objects, seeking to compare neural correlates of learning at the levels of individual visual objects and of higher-order temporal dependencies. We focus on the visual pathways in the ventral occipitotemporal cortex, the major neural substrate of visual experience and long-term memory (reviewed by Grill-Spector and Weiner 2014; Bi et al. 2016; Kravitz et al. 2013; Weiner and Zilles 2016). We hypothesize that learning of visual shapes (*i.e.*, spatiotemporal relationships of characteristic features) might interact with the learning of the context in which such shapes appear (*i.e.*, spatiotemporal configurations of distinct shapes). By mapping the influence of context on shape representations, we hope to improve our understanding of statistical learning at distinct timescales. Insofar as both invariant shape and higher-order temporal context are aspects of the causal structure of the visual environment, our findings may also pertain to the neural substrates of “structural learning” (Tervo et al., 2016).

Our visual stimuli are synthetic, three-dimensional objects of unique and characteristic shape, which are presented many times over several days, but every time in a different way (from different points of view and rotating about different axes). To locate neural representations of object shapes that are invariant to these differences, we monitor multivariate BOLD activity as observers gain familiarity with a particular set of objects. Specifically, we assess the ‘representational similarity’ (Kriegeskorte et al., 2008 a; Haxby, 2012) of activity evoked by repeated presentations of the same objects and of different objects. A companion study reports how the representational geometry of “object identity” changes with familiarity (Kakaei and Braun, 2023).

To assess how shape representations are modified by temporal context, we follow Schapiro and colleagues (Schapiro et al., 2013) and manipulate the order in which objects are presented, such that objects either form clusters with pairwise sequential dependencies (“temporal communities”) or avoid such dependencies. Such “temporal communities” are behaviourally relevant and influence the order in which different objects become familiar (Kakaei et al., 2021). After discounting the effects of temporal proximity, we assess the extent to which neural representations of “temporal communities” (lasting approximately 30 s) are superimposed on the neural representations of “object identity” (viewed for 3 s). In short, we compare correlates of statistical learning at two different time scales.

Our results revealed representations of temporal communities over much of the ventral occipitotemporal cortex, at similar levels of the visual hierarchy as representations of object identity. Approximately 20% of the identity-selective sites were also community-selective. In many cases, objects marking transitions between communities were represented distinctly from other objects. A few sites in the frontal cortex and anterior temporal cortex exhibited community selectivity without significant identity selectivity. We conclude that at the sensitivity afforded by fMRI, hierarchical levels of representation of visual object identity and of regularities in their order of presentation appear to be largely coextensive. Thus, recognition learning and “structure learning” may be subserved by overlapping cortical networks.

## 2 Methods

The experimental paradigm and procedure are described in detail elsewhere (Kakaei and Braun, 2023). Here we only summarize the most pertinent aspects.

### 2.1 Participants

Eight healthy participants (4 female and 4 male; aged 25 to 32 years) took part in the functional imaging experiment. Twelve additional participants performed in a purely behavioral experiment (Kakaei et al., 2021).

### 2.2 Experimental Paradigm

Synthetic visual objects were generated and presented as described previously (Kakaei et al., 2021; Kakaei and Braun, 2023). Objects were presented 3 *s* from varying points of view while rotating slowly around variously oriented axes in the frontal plane (**Fig. 1A**). Fifteen objects recurred at least 190 (mean±S.D: 216 ± 9) times each over three sessions (‘recurring’ objects), whereas 360 other objects appeared exactly once (‘non-recurring’ objects). Observers responded to each object by classifying it as either ‘familiar’ or ‘unfamiliar’. Over the course of three sessions, all observers gradually became familiar with the ‘recurring objects’ (**Fig. 1C**).

**Figure 1:**
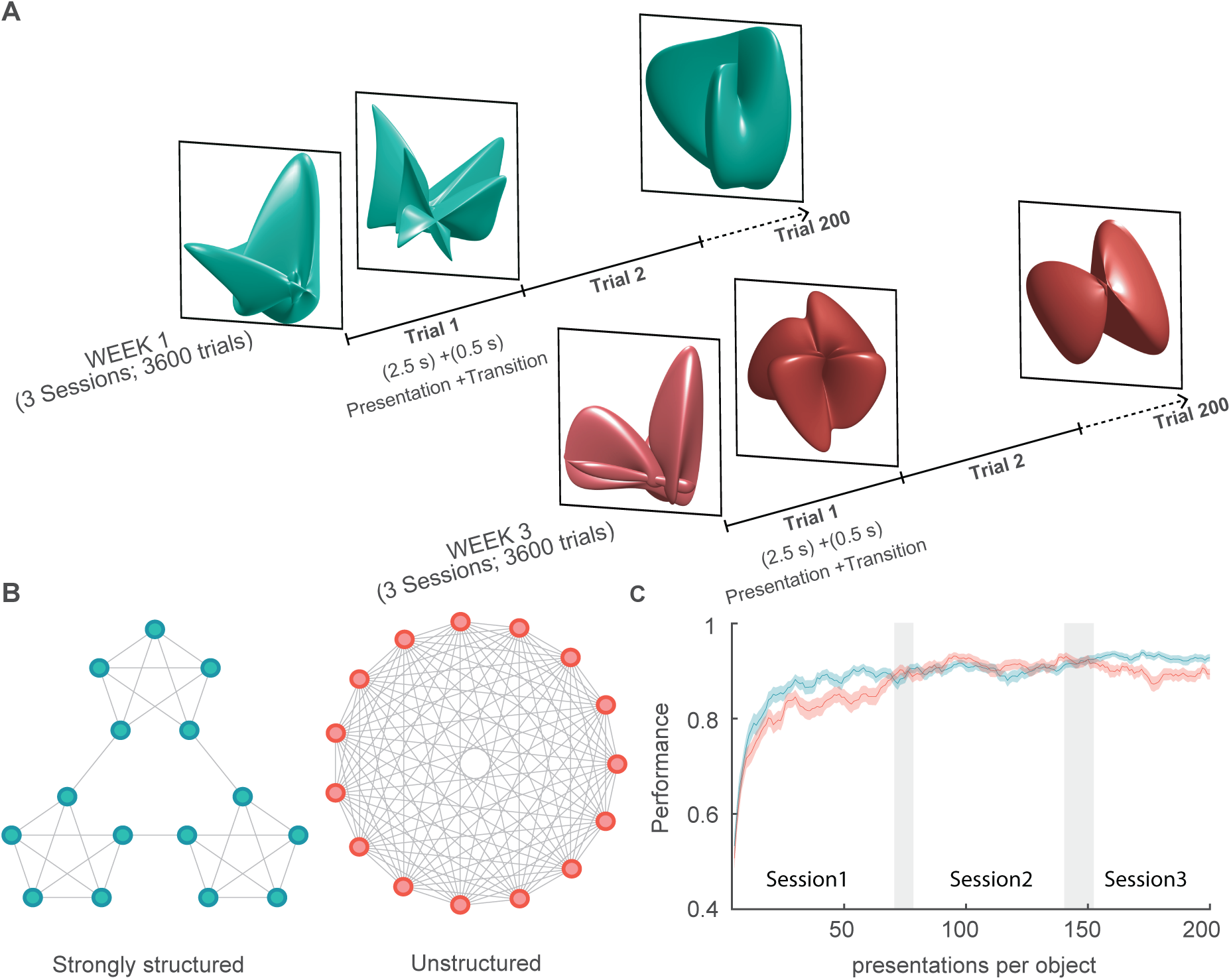
Experimental paradigm. **A)** Observers viewed complex three-dimensional objects, slowly rotating, in sequences of 200 presentations (2.5s presentation time and 0.5s transition time). Most objects (180 of 200) recurred multiple times within and between sequences (‘recurring’ objects). The others (20 of 200) were presented only once (‘non-recurring’ objects). Observers viewed 18 sequences in three sessions and attempted to classify each object as either ‘familiar’ or ‘unfamiliar’. **B)** Sequences were generated from quasi-random walks on either a sparse & modular graph, or a fully-connected & non-modular graph (nodes are recurring objects, and links are possible successions). Sequences from the modular graph (left) exhibit clustered sequential dependencies and are ‘strongly structured’. Sequences from the fully-connected graph (right) have no such dependencies and are ‘unstructured’. **C)** Over three sessions, observers learned to classify recurring and non-recurring objects respectively as ‘familiar’ and ‘unfamiliar’. Adapted from Kakaei et al. 2021.

Presentation sequences were generated as quasi-random walks on graphs that represented 15 recurring objects as nodes and possible continuations as edges (**Fig. 1B**; Kakaei et al. 2021). Each sequence began at a random node and continued with equal probability on any one of the available edges, except that immediate repetition (*X* → *X*) and direct returns (*X* → *Y* → *X*) were not allowed. Each sequence comprised 200 objects and lasted 600 *s*. Most objects (180 of 200) were recurring, with each individual object being repeated 12 ± 1.9 times on average. Non-recurring objects (20 of 200) were interspersed at random intervals.

Strongly structured sequences were generated from the modular graph depicted left in **Fig. 1B**. In this graph, each object is linked to exactly four other objects (*i.e.*, may be preceded or followed by four other objects). However, these links are clustered such as to form three “communities” of five objects each. When sequences are generated from this graph, approximately 9 ± 2 successive objects derive from the same community.

The 105 possible pairings of 15 objects may be divided into four groups, as illustrated in **Fig. 5**. 27 pairs are from the same community and also adjacent on the graph (SA pairs). 3 pairs are from different communities, but adjacent on the graph (DA pairs), whereas further 3 pairs are from the same community, but non-adjacent on the graph (SN pairs). Finally, 72 pairs are from different communities and non-adjacent on the graph (DN pairs).

Note that only *SA* pairs and *DA* pairs are possible in strongly-structured sequences, in the sense that one member directly follows the other. All possible pairs are equally likely to occur (probability approximately 1*/*60), even though there are fewer *DA* pairs (3 pairs) than *SA* pairs (27 pairs) (for details see Kakaei et al. 2021).

Unstructured sequences were generated from the graph depicted right in **Fig. 1B**. In this graph, each object is linked to all other objects (*i.e.*, it may be preceded or followed by any one of the other objects), so that no sequential dependencies arise.

In the first week of the experiment, participants experienced one set of fifteen recurring objects in one type of sequence whereas, in the third week, they experienced another set of fifteen objects in the other type of sequence. In each week, participants performed in three fMRI sessions (on three separate days), each time viewing six sequences with 200 objects. The second week of the experiment was free.

### 2.3 fMRI data analysis

To study the effect of sequence structure on the neural representation of object shape, we extracted the multivoxel activity pattern at *N_t_*=9 time points following object onset. In a functional parcel with *N_vox_* voxels, this response pattern constituted a point (or vector) in a *N_dim_* -dimensional space, where *N_dim_* = *N_t_* · *N_vox_* (**Fig. 2.A**). Our objective was to compare distances between response patterns to the same objects, to different objects in the same community, and to different objects in different communities, in other words, to analyse representational similarity or dissimilarity in terms of the standardized Euclidean (Mahalonobis) distance between responses in a high-dimensional space (Kriegeskorte and Diedrichsen, 2019). Over all 758 parcels, the mean and standard deviation of the response dimensionality was *N_dim_* = 1911 ± 634, with a range from 405 to 4158.

**Figure 2:**
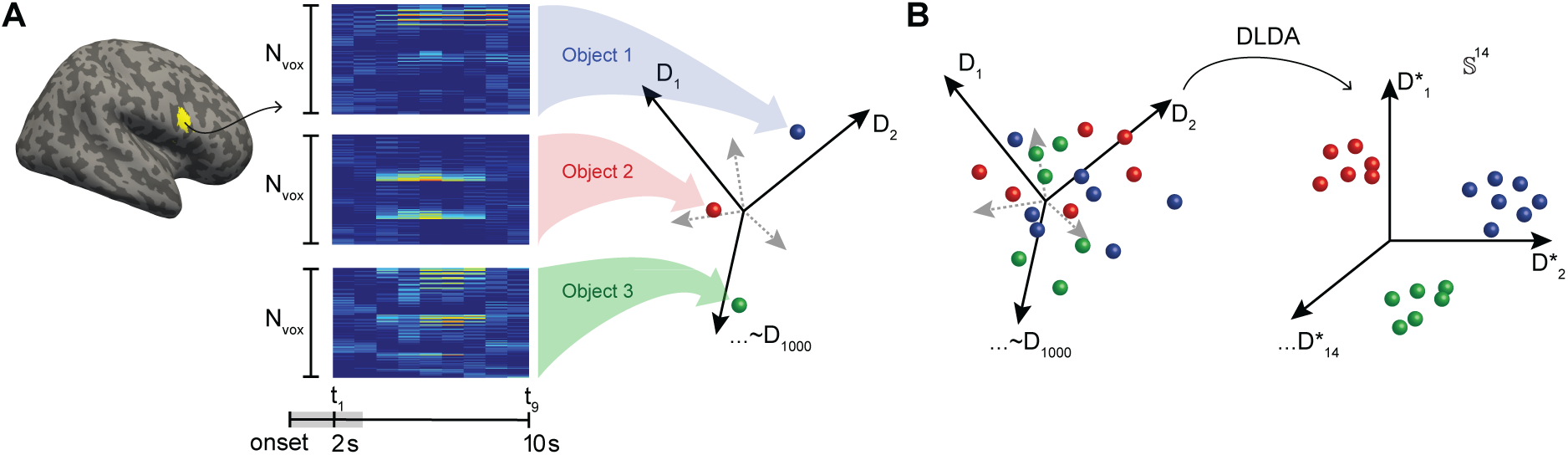
Direct linear discriminant analysis (DLDA) of multivariate BOLD signals. For each functional parcel, we identified a 14-dimensional space that optimally discriminated the 15 classes of activity patterns associated with *recurring* objects. **A)** For a given parcel with *Nvox* voxels (here in yellow: Frontal-Inf-R-8), activity was recorded over 9 *s* during and after object presentation (from 2 to 11 *s* after onset). Each such activity pattern corresponds to a point in a 9 · *Nvox* -dimensional vector space (right), here represented schematically by spheres (red, green, and blue). **B)** In the optimally discriminative subspace S ∈ S^14^, the representational similarity of multiple responses to the same objects could be established. Adapted from Kakaei and Braun 2023.

To focus the analysis on directions of variance pertinent to our objective, we reduced dimensionality with Fisher’s Linear Discriminant Analysis (LDA) for multiple classes to identify the (at most) (*κ* − 1)-dimensional subspace S that optimally discriminates *κ*=15 classes of activity patterns. Optimality is defined here as simultaneously minimizing within-class variance and maximizing the between-class variance of activity patterns. This approach corresponds to a ‘supervised’ principal component analysis and yields (*κ*−1) comparatively informative dimensions. For example, of the total response variance of one particular parcel (405 Occipital-Mid-L, with 194 voxels and *N_dim_* =1746), a share of 38% was captured by the optimally discriminative 14-dimensional space S, whereas a share of 59% was captured by the first 14 principal components. However, the variance captured by subspace S was distributed more uniformly over dimensions (3 ± 3% per dimension) than in the principal component subspace (4 ± 6% per dimension).

We analyzed the representation of sequence structure with data from strongly structured sequences (8 observers), performing identical analysis on data from unstructured sequences (8 observers) for comparison. As detailed further below, spurious ‘effects’ of community structure can be observed due to systematic and/or unsystematic fluctuations of responsiveness over time. To guard against such spurious effects, we removed the effects of temporal proximity and verified that our analyses yielded null results with data from unstructured sequences.

#### 2.3.1 Linear discriminant analysis

A numerically tractable procedure for identifying the optimal subspace S is available in terms of ‘direct LDA’ or DLDA (Yu and Yang, 2001; Ye et al., 2006). Briefly, this method first diagonalizes between-class variance to identify *κ* − 1 discriminative eigenvectors with non-zero eigenvalues, next diagonalizes within-class variance and finally yields a rectangular matrix for projecting activity patterns from the original activity space (dimensionality *N_dim_*) to the maximally discriminative subspace S and back. As this method is linear and relies on all available degrees of freedom, its results are deterministic. Specifically, it results in dimensionally reduced activity patterns *x_jk_*, where *j* ∈ {1*, . . .,* 14} indexes (reduced) dimensions and *k* indexes trials. The link github.com/cognitive-biology/DLDA provides a Matlab implementation of DLDA.

#### 2.3.2 Amplitudes, distances, and correlations

Activity patterns *x_jk_* associated with trials *k* were analyzed in the maximally discriminative subspace S. The average normalized amplitude 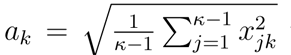 was ⟨*a*⟩ = 0.99 and the average normalized distance 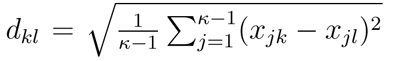 between patterns from trials *k* and *l* was ⟨*d*⟩ = 1.40, consistent with the expected distance between random patterns 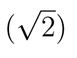

The pattern associated with successive trials exhibited a weak temporal correlation, with approximately 5% smaller distances at delays below 4 trials and approximately 2% larger distances at delays ranging from 6 to 15 trials. **Supplementary Figure S2** shows the delay-dependent distance between response pairs, as well as the pairwise distance within runs, averaged over all parcels and observers.

To discount this temporal correlation (Section 2.3.3), we established for each parcel *w* the average delay-dependent distance *T_w_*(Δ*i*) = ⟨*d_w,u,r_*(Δ*i*)⟩*_u,r_* between patterns with relative delay Δ*i*, where the average was taken over subjects *u* and runs *r*. The timecourse *T_w_* allowed us to subtract the average effect of temporal correlation by computing residual distance *d^corrected^* (Δ*i*) = *d_w,u,r_*(Δ*i*) − *T_w_*(Δ*i*) + ⟨*T_w_*(Δ*i*)⟩_Δ_*_i_*, where ⟨*T_w_*(Δ*i*)⟩_Δ_*_i_* is the average value over delays.

#### 2.3.3 Geometry of temporal community representations

To assess the extent to which community structure was reflected in the neural representation of objects, we compared pairwise distances between responses to objects within and between communities. Specifically, we first obtained pairwise distances *d_ij_* and sorted them into two groups: within-community distances with average distance = ⟨*d_ij_*(*ij*|*i, j* ∈ *L*)⟩ and between-community distances with average distance = ⟨*d_ij_*(*ij*|*i* ∈ *L, j* ∈ *K, L* = *K*)⟩. Then, we established the signed difference Δ*^BW^* = ⟨*D^between^*⟩ − ⟨*D^within^*⟩ and assessed the statistical significance of Δ*^BW^* with a two-sample t-test. After correcting for false discovery (Benjamini and Hochberg, 1995), we summarized the results for each parcel in terms of *t* -statistics *t^BW^* .

A similar procedure was used to assess differences between classes of object pairs. Specifically, for every parcel, we established the average pairwise distance *D_w_* (averaged over all pairs and all observers) for different classes of object pairs: adjacent & same community (SA), non-adjacent & same community (SN), adjacent & different communities (DA), and non-adjacent & different communities (DN). The resulting values were termed *D^SA^*, *D^DA^*, *D^SN^* and *D^DN^* . The statistical significance was assessed by comparing the observed values to the pair-wise distance *D^diff^*, which contains pair-wise distances of all 4 types of object pairs, by a two-sampled *t*-test and the results were summarized in terms of z-scores *t_w_^SA^*, *t_w_^DA^*, *t_w_^SN^* and *t_w_^DN^* .

Note that response distances within and between communities are confounded by temporal proximity because responses *within* communities tend to have shorter relative latencies than responses *between* communities (Schapiro et al., 2013). To assess the degree to which temporal proximity contaminates the observed community signal, we repeated the analysis of community representations for different ranges of temporal latencies. Specifically, we recalculated the average pairwise distances *D^between^* and *D^within^*, and the corresponding *t^BW^* for object pairs *i, j* whose relative latencies *τ_ij_* where bounded from below by *τ_LB_* ⩽ *τ_ij_* and from above by sequence termination, with the lower bound ranging over *τ_LB_* ∈ {1*, . . .,* 30}. The *t*-statistics of response pairs with bounded latencies and their corresponding p-values, corrected for false discovery rate, will be denoted as *t^BW^* (*τ_LB_*) and *P^BW^* (*τ_LB_*), respectively.

To assess whether community representations are consistent over different latency ranges, we examined how *t^BW^* (*τ_LB_*) changes with its lower bound *τ_LB_* . Specifically, for each parcel, we defined a consistency measure *τ_sig_* as the highest lower bound at which *t^BW^* (*τ_sig_*) remains significant. We considered a parcel as *‘community selective’* only if *τ_sig_* ≥ 30. In other words, a ‘community selective’ parcel exhibited significant between-community separability *t^BW^* for all lower bounds *τ_LB_* ∈ {1*, . . .,* 30}. This ruled out the possibility that community selectivity was a spurious effect of temporal proximity (which was strongest at shorter latencies).

#### 2.3.4 Statistical power

The representational similarity analysis with respect to object identity described in Kakaei and Braun 2023 was based on approximately 216 object responses (18 sequences with approximately 12 recurrences of each object) from each of 16 observers, affording approximately 370, 000 representational distances for each of the 105 object pairs. In contrast, the assessment of representational similarity with respect to community was based on approximately 120 community episodes (18 sequences with approximately 6 recurrences of each community) from each of 8 observers, affording approximately 57, 000 representational distances for each of the 3 community pairs. Hence, the number of independent pairwise observations pertaining to identity was approximately 225 times larger than the number pertaining to community. Accordingly, on purely statistical grounds, the sensitivity of our paradigm for detecting community selectivity is expected to be approximately 15 times *lower* than for detecting identity selectivity.

#### 2.3.5 Dimensional reduction

To visualize the representational geometry of object shape and community structure, we calculated a distance matrix *D_w,u,r_*(*i, j*) = ⟨*d_ij_*⟩ of response distances corrected for temporal proximity within each run *r*, for every parcel *w* and observer *u*. Averaging over the runs produced matrices *D_w,u_* of size 15 × 15 average distances in the discriminative subspace S.

As both presentation sequences and neural representations of object shape differed between observers, these matrices could not be averaged directly over observers. To sidestep the difficulty, we permuted the object order of the matrix 10^4^ times while maintaining graph structure (adjacency and module membership), to first obtain an ensemble average matrix 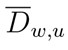 for each observer, and finally the observer average 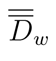 of ensemble averages.

Using multidimensional scaling (Matlab function *mdscale*), we converted the observer average matrix 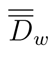 to a two-dimensional map of 15 locations approximating these pairwise distances. Note that all maps exhibit a three-fold rotational symmetry owed to the permutation procedure.

## 3 Results

### 3.1 Representation of temporal community structure

To assess the extent to the which temporal community structure of presentation sequences was reflected in neural representations, we compared pairwise distances of responses within communities and between communities, in the maximally discriminating space. As illustrated in **Fig. 3A**, pairwise distances were computed for all observers, runs, and parcels *w* (including identity-selective parcels), to obtain average pairwise distance *D^within^* communities, average pairwise distance *D^between^* between communities, and the average separability Δ*^BW^* = *D^between^* − *D^within^*. For every parcel *w*, we assessed whether or not Δ*^BW^* values differed significantly from zero with t-statistic *t^BW^* . As this analysis was heavily confounded by temporal auto-correlations, we carried out two corrections intended to help dissociate community selectivity and temporal proximity (see section 2.3.2). Firstly, for each raw pairwise distance and its latency, we computed a residual pairwise distance by subtracting the average distance at that latency. Secondly, we compared the t-statistic *t^BW^* for subsets of pairwise distances covering different temporal latency ranges (*τ_LB_*≤ *τ* ≤ 30; *τ_LB_*∈ {1*, . . .,* 30}). For all parcels, significance decreased monotonically when lower bound *τ_LB_* was raised and shorter latencies were progressively excluded. Thus, the situation was summarized by the *largest* value of *τ_LB_* at which *t^BW^* statistic was significant, which value was termed *τ_sig_* . A high value of *τ_sig_*indicated significance over all latency ranges, both including shorter latencies (low values of *τ_LB_*) and excluding shorter latencies (high values of *τ_LB_*). A low value of *τ_sig_* indicated significance only for ranges that included shorter latencies (low values of *τ_LB_*).

**Figure 3:**
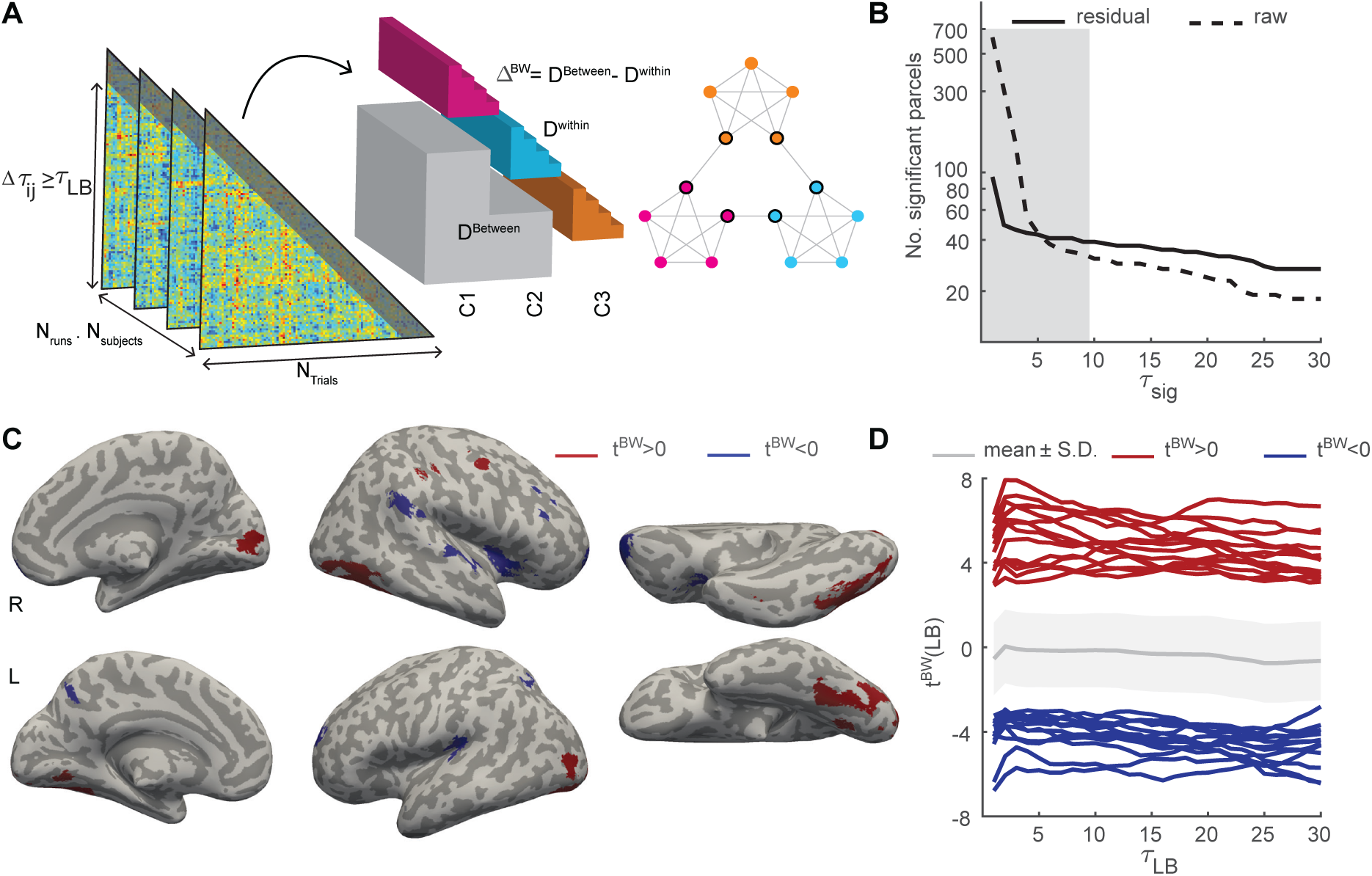
Neural representation of temporal community structure. **A)** Pair-wise distances between object responses (triangular matrices) were corrected for the average auto-correlation, thresholded by latency *τ_ij_* (lower bound *τ_LB_*, indicated by shading) and sorted into different subsets – within-community pairs (cyan, magenta, orange) and between-community pairs (grey) – according to object positions on the modular path (right). For the average signed difference Δ*^BW^*, statistic *t^BW^* was computed. **B)** Number of parcels with consistent significance up to *τ_sig_* for residual (solid) and raw (dashed) pairwise distances. The average duration of a community visit was 9.4 ± 0.15 (gray shading). **C)** Representation of ‘temporal communities’ by parcels with consistently significant Δ*^BW^* . In 14 parcels (red), between-community pairs are significantly more separable (*t^BW^ >* 0, corrected *p <* 0.05) over all latency ranges whereas, in 13 parcels (blue), within-community pairs are more separable (*t^BW^ <* 0) over all ranges. **D)** Between-community separability *t^BW^* for different latency ranges (lower bound *t^BW^*), for ‘community-selective’ parcels with positive *t^BW^ >* 0 (red) and negative *t^BW^ <* 0 (blue). The mean and S.D. of separability over all parcels are shown in gray.

When raw pairwise distances were used, almost all parcels (613 out of 758 parcels) exhibited significant separability Δ*^BW^* . When residual pairwise distances were considered, ninety-three parcels retained significant Δ*^BW^* (left margin of **Fig. 3B**). In 28 of these 93 parcels, between-community separability was higher (*t^BW^ >* 0) and in the remaining parcels, it was lower (*t^BW^ <* 0).

This disparity between raw and residual distances shows that community structure is confounded by temporal auto-correlation to a considerable degree. This is also evident from strong dependence of *t^BW^* on the range of temporal latencies *τ_sig_* (**Fig. 3B**). When only latency ranges including shorter latencies are considered (*τ_sig_*≤ 5) many more parcels are consistently significant than when ranges excluding shorter latencies are also considered (*τ_sig_ >* 15). Applying the strictest criterion and considering only parcels with significant Δ*^BW^* (FDR corrected *p <* 0.05) for all latency bounds *τ_LB_* ∈ {1*, . . .,* 30} (*τ_sig_* = 30), we obtained 27 parcels that we considered *‘community selective’*. These parcels are listed in **Table A.1** and illustrated in **Fig. 3CD** and in **Supplementary Fig. S1**.

The above analysis yielded interpretable results for strongly-structured presentation sequences, where every object can be objectively assigned to one particular community. When the analysis was repeated for unstructured presentation sequences (by counterfactually assuming a structured sequence and assigning communities accordingly), no systematic results were obtained, as shown in **Supplementary Fig. S3**. Specifically, apparent community selectivity is observed only when uncorrected distances over low-latency ranges are considered. Correcting for temporal correlations eliminates this spurious selectivity. The static matrix of average pairwise distances provides an instructive baseline for spurious community selectivity that is entirely due to temporal correlations. Apart from very short latencies, the results from this matrix are comparable to results from unstructured sequences, both for *positively* and *negatively* community selective parcels temporal correlations (**Supplementary Fig. S3BC**). Results for structured sequences are dramatically different (both higher and lower), corroborating the validity of our analysis of community selectivity.

Fourteen community-selective parcels with *higher* separability of between-community pairs (Δ*^BW^ >* 0) were located in occipital and inferior-temporal regions, as well as in frontal and postcentral cortex. Eleven of these parcels were also identity-selective. Thirteen other parcels exhibited significantly *lower* separability of between-community pairs (Δ*^BW^ <* 0) and were located in the superior-, middle- and orbitofrontal cortex, as well as in insular, superior parietal, supramarginal, and superior temporal cortex. Only one of these parcels was also identity-selective.

The respective cortical distributions of the representations of object identity and community membership are compared and illustrated in **Fig. 4**. The criterion for community-selectivity was a significantly positive or negative *t* -score value *t^BW^*, whereas the criterion for identity-selectivity was a significantly positive minimum statistics of classification accuracy *α_min_* (for details, see Kakaei and Braun 2023). **Fig. 4C** shows average classification accuracy *α^identity^* as well as *α_min_* . Coloring indicates whether parcels combined identity-selectivity with *positive* community-selectivity (11 parcels, orange) or *negative* community-selectivity (1 parcel, cyan), or whether parcels were either exclusively community selective (3 parcels *positively* in red, 12 parcels *negatively* in blue) or exclusively identity-selective (112 parcels, yellow). Of the 124 identity-selective parcels, 12 parcels (approximately 10%) were additionally community-selective. Jointly selective parcels were most common in the mid-level visual cortex (ventral occipital cortex, lingual and fusiform gyrus) and somewhat less common in the early visual cortex (V1, V2, V3, hV4). Jointly selective parcels were largely absent from high-level visual areas in the parietal and frontal cortex (inferior parietal sulcus, superior parietal lobule, insula, inferior and medial frontal cortex), but were present in the anterior inferior temporal cortex. The one negatively community-selective parcel in the intraparietal sulcus appeared to be an exception. In summary, jointly selective parcels were present at all levels of the ventral visual pathway.

**Figure 4:**
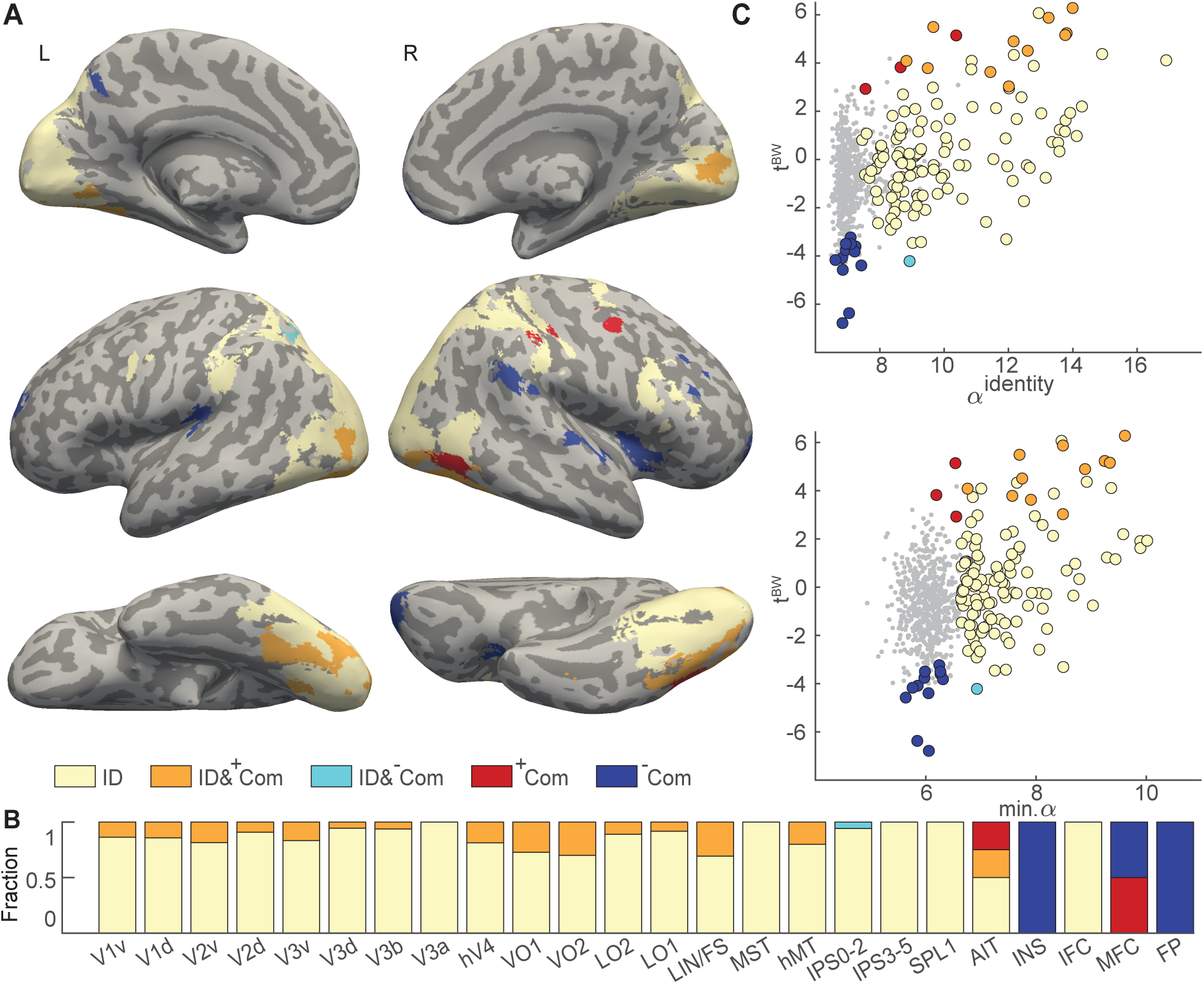
Object identity and community membership compared. **A)** Anatomical distribution of 142 parcels that are identity-selective, community-selective, or both. 11 parcels (orange) are both identity-selective and *positively* community- selective, while 1 parcel (cyan) combines identity-selectivity with *negative* community-selectivity. 15 parcels are exclusively community-selective, 3 parcels *positively* (red) and 12 parcels *negatively* (blue). The remaining 112 parcels (yellow) are exclusively identity-selective. **B)** Share of identity and community representation in 144 parcels with significant representation, assigned to 29 topographical regions. Coloring corresponds to **A** and indicates the fraction of voxels from parcels with different selectivity. Visual cortex (V1-hV4), ventral occipital cortex (VO), lateral occipital cortex (LO), lingual and fusiform gyri (LIN/FS), medial temporal areas (MST, hMT), intraparietal sculcus (IPS), superior parietal lobule (SPL), anterior inferior temporal cortex (AIT), insula (INS), inferior frontal cortex (IFC), medial frontal cortex (MFG), and frontal pole (FP). **C)** Quantitative comparison of selectivity for identity and community over all parcels. Identity-selectivity is quantified either by average classification accuracy *α^identity^* (top) or by the minimum statistic of classification accuracy (bottom). Community selectivity is measured by positive or negative values of *t^BW^* . Significantly selective parcels are represented by colored discs, and non-selective parcels by grey dots. Coloring corresponds to **A**.

Note that the comparison of community- and identity selectivity was skewed by disparate statistical power. The assessment of community selectivity was based on approximately 225 times fewer observed response distances than the assessment of identity selectivity (see Methods) so statistical sensitivity was expected to be approximately 15 times lower. Accordingly, if community selectivity was detected in only a fraction of identity-selective parcels, this could in part have been due to lower statistical power.

Nominally non-identity-selective parcels with *positive* community-selectivity were located in the anterior inferior temporal cortex and in the medial frontal cortex. As seen in the top panel of **Fig. 4C**, the average classification accuracy *α^identity^* of these 3 parcels was comparable to other identity-selective parcels. However, these parcels just missed the minimum statistics criterion for significance, as seen in the bottom panel. It seems possible that community selectivity degraded identity selectivity in these parcels, in the sense that reduced response distances within a community might also have reduced distances between the different objects of this community.

Non-identity-selective parcels with *negative* community-selectivity were located in the insula, the medial frontal cortex, and at the frontal pole. These 11 parcels exhibited no trace of identity selectivity in terms of either the observer average or the minimum statistics. Negative selectivity implies that responses to objects from different communities were more similar than responses to objects from the same community. As discussed below, it seems possible that the responses in these areas placed particular emphasis on ‘linking objects’, thereby highlighting the ‘novelty’ or ‘surprise’ associated with the transition to another community and the appearance of unexpected objects.

Alternatively, it is possible that responses in these areas are simply encoding the dimensions of the task-space (familiar ↔ novel) in a kind of ‘cognitive-map’ Behrens et al. (2018). Recall that observers were asked merely to classify objects as ‘familiar’ or ‘novel’ (rather than to identify them ‘by name’) so that there was no behavioral benefit to maintaining individual object representations.

### 3.2 Representation of object pairs

In addition to temporal communities, a prominent feature of strongly structured sequences was the ‘linking objects’ that marked transitions out of the previous community and into the next community. Accordingly, we wondered to what extent the linking objects contributed to the representation of temporal communities. To address this issue, we compared the overall measure *t^BW^* with the pairwise separability of specific types of object pairs: non-adjacent objects in different communities (DN), non-adjacent objects in the same community (SN), adjacent objects in different communities (DA), and adjacent objects in the same community (SA). The results are shown in **Fig. 5**. The separability measure *t^SA^* was negatively correlated with *t^BW^* (*ρ* = −0.91, *p <* 0.01), whereas the measure *t^DN^* was positively correlated with *t^BW^*(*ρ* = 0.93, *p* ≪ 0.01). The separability measures *t^DA^* and *t^SN^*were also negative correlated with *t^BW^*, though much less strongly (*ρ* = −0.15, *p <* 0.01 and *ρ* = −0.19, *p <* 0.01; respectively). These results were robust and held for all lower temporal bounds *τ_LB_* ≤ 30, except for the correlation between *t^BW^* and *t^DA^*, which held only for *τ_LB_* ≤ 28.

These results show that the representation of community structure (indexed by *t^BW^*) includes a reduced separation of *SA* pairs (indexed by *t^SA^*), as well as an increased separation of *DN* pairs (indexed by *t^DN^*). Recall that *SA* (and *DA*) pairs occur in presentation sequences (with probability 1*/*60), whereas *SN* (and *DN*) pairs never occur. The selective modulation of representational distance for one of the two adjacent (and therefore occurring) pairs, appears to be a correlate of temporal community structure. The same can be said for the selective modulation of representational distance for one of the two non-adjacent (and therefore non-occurring) pairs. Furthermore, the correlation between community representation and separation of *SA* and *DN* pairs is evident not only in the few parcels meeting the statistical threshold for community selectivity (red and blue dots in **Fig. 5**), but in all other parcels as well (grey dots in **Fig. 5**). Thus, reduced separation of *SA* pairs and increased separation of *DN* pairs appear to be a general correlate of community structure within the cortical representation of object identity.

The results described above depend critically on the correction for temporal correlations (**Supplementary Fig. S4**). Without this correction, the *t^BW^* measure for between-community separation is dominated by the influence of short-latency pairs. When shorter latencies are excluded and *τ_LB_* ≥ 5, the correction ceases to make a difference. This underlines again that correcting for average temporal correlations is key to establishing representations of community structure.

### 3.3 Representational space

A previous study with structured sequences (Schapiro et al., 2013) reported that within-community distances are typically smaller than between-community distances and elegantly illustrated this with multidimensional scaling. We sought to replicate this finding by visualizing the differential proximity of different object pairs in the response space of individual parcels. To obtain interpretable results, we employed a permutation procedure that allowed us to average proximity matrices over observers (see Methods for details). The resulting illustrations exhibit a three-fold rotational symmetry that is artificial and owed to this permutation procedure.

For the fourteen *positively* community-selective parcels, the differential proximity of different object pairs is illustrated in (**Fig. 6**). In all cases except one, objects were clustered by community (*i.e.*, spaced more closely within than between communites), with Temporal-Inf-R-for providing the most extreme example. Additionally, ‘linking’ objects tended to be positioned differently than internal objects, in all but two cases closer to each other (and to the center) (Calarine-L-9, Calcalrine-L-5, Lingual-L-1, Occiptal-Mid- L-4, Occiptial-Mid-L-9, Occipital-Inf-L-2, Fusiform-L-2, Fusiform-L-6, Postcentral-R-11, Temporal-Inf-R-10). Exceptions were Frontal-Mid-R-7, where only internal objects clustered by community, and Occipital-Inf-R2/4, where linking objects were more distant from each other. As these illustrations reflect only relative distances, **Supplementary Fig. S6** shows absolute distances in terms of the average and standard deviation over parcels, separately for internal objects and linking objects, as well as within and between communities. Both effects mentioned here – clustering by community and relative proximity of linking objects – were statistically significant.

**Figure 5:**
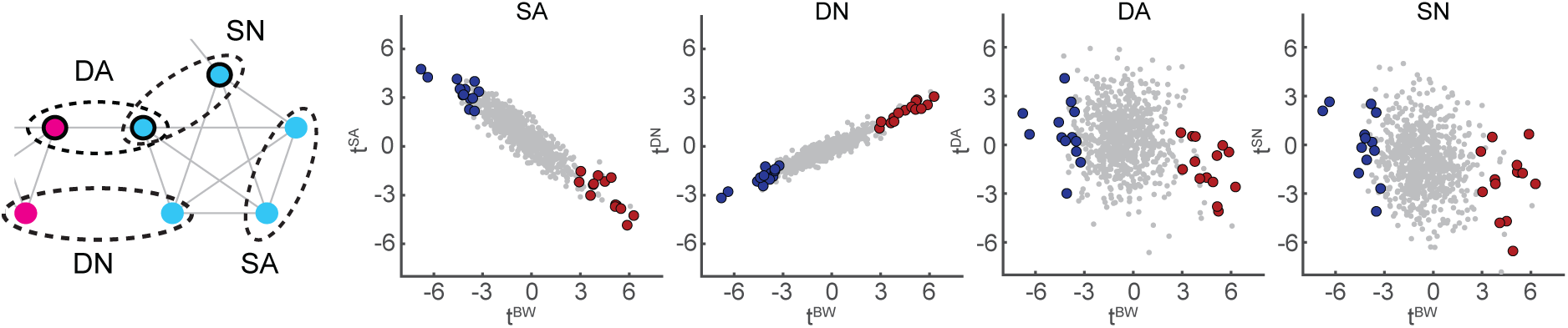
Neural representation of detailed aspects of sequence structure: same (*S*) or different (*D*) community, and adjacent (*A*) or non-adjacent (*N*) position on the path. Differential separability of between- and within-community pairs are represented by t-score value *t^BW^* and compared to separability *t^SA^* of adjacent objects in the same community (left), *t^DN^* of non-adjacent objects in different communities (middle left), *t^DA^* of adjacent objects in different communities (middle right), and *t^SN^* of non-adjacent objects in the same community (right), for all 758 parcels. Community-selective parcels are shown in red or blue (compare **Fig.3C**).

**Figure 6:**
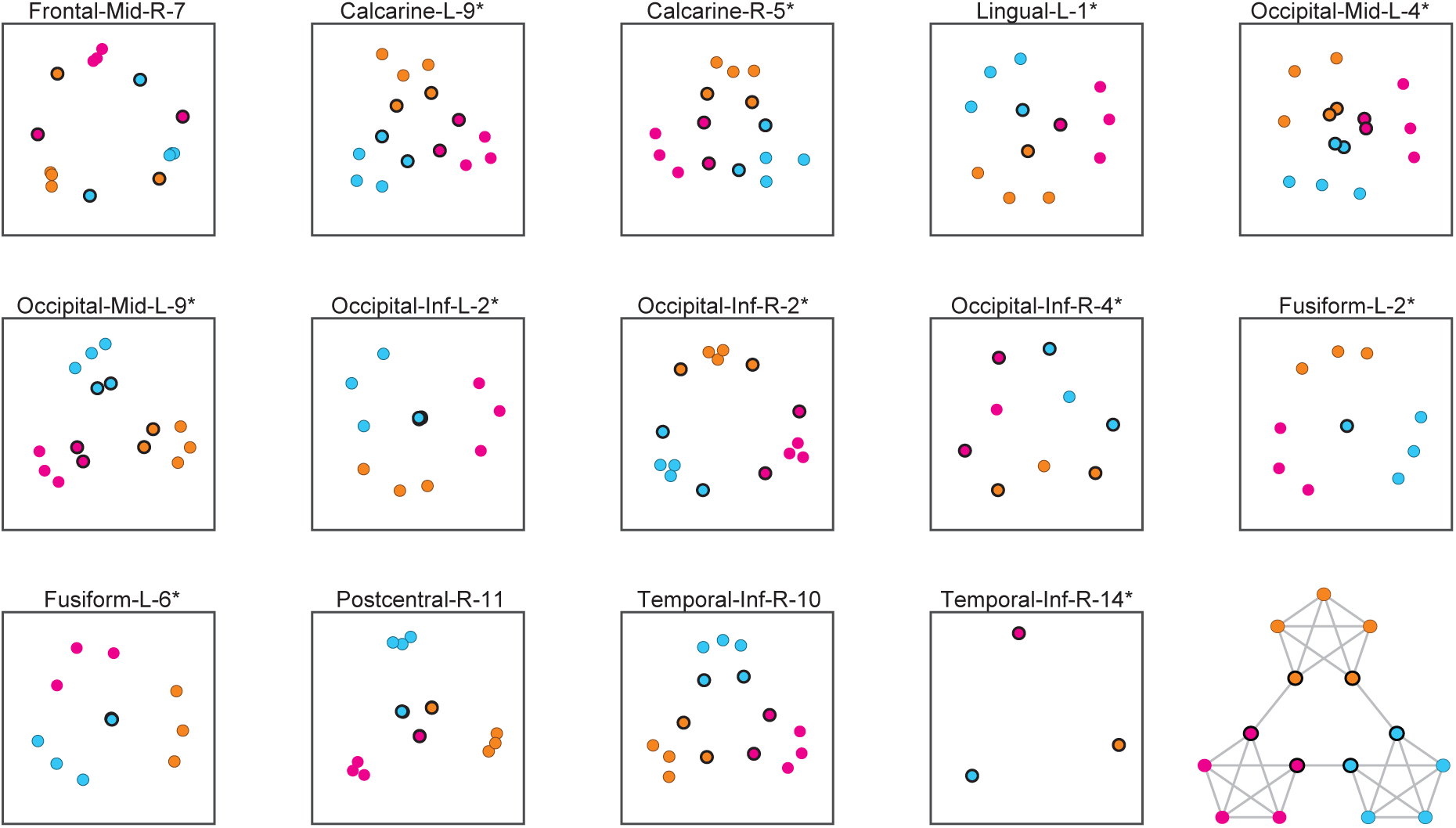
Representation of temporal community structure in positively selective parcels. Multidimensional reduction of the pair-wise distance matrix averaged over path permutations and over observers. Communities are distinguished by color and linking objects by a black outline, as indicated by the path diagram (inset). Fourteen parcels exhibited higher separability between communities than within communities (*t^BW^ >* 0). Identity-selective parcels are marked with *\**.

Results for the thirteen *negatively* community-selective parcels are shown in **Fig. 7**. The clustering of internal objects (Frontal-Sup-L-12, Frontal-Sup-Orb-R-3, Frontal-Mid- R-16, Frontal-Med-Orb-R-3, Parietal-Sup-L-8, and Temporal-Sup-R-6) was variable but, when averaged over parcels, internal objects were more distant within than between communities (**Supplementary Fig. S6**). Linking objects were often distant from each other (Frontal-Sup-R-19, Frontal-Sup-Orb-R-3, Insula-R-6, Parietal-Sup-L-8, Precuneus-L-12, Putamen-R-6) and, on average, closer to internal objects of different communities than of the same community. In other words, the representation emphasized the linking objects associated with a transition between communities. In six parcels, all linking objects were distant from each other (and from the center), suggesting that the representation in these parcels individuated different transitions between communities (Frontal-Sup-R- 19, Frontal-Sup-Orb-R-3, Insula-R-6, Parietal-Sup-L-8, Precuneus-L-12, Putamen-R-6). However, in seven other parcels, linking objects were separated less well than internal objects (Frontal-Sup-L-12, Frontal-Mid-R-16, Rolandic-Oper-L-2, Frontal-Med-Orb-R-3, Insula-R-5, SupraMarginal-R-4, Temporal-Sup-R-6), suggesting that the representation conflated different transitions between communities.

**Figure 7:**
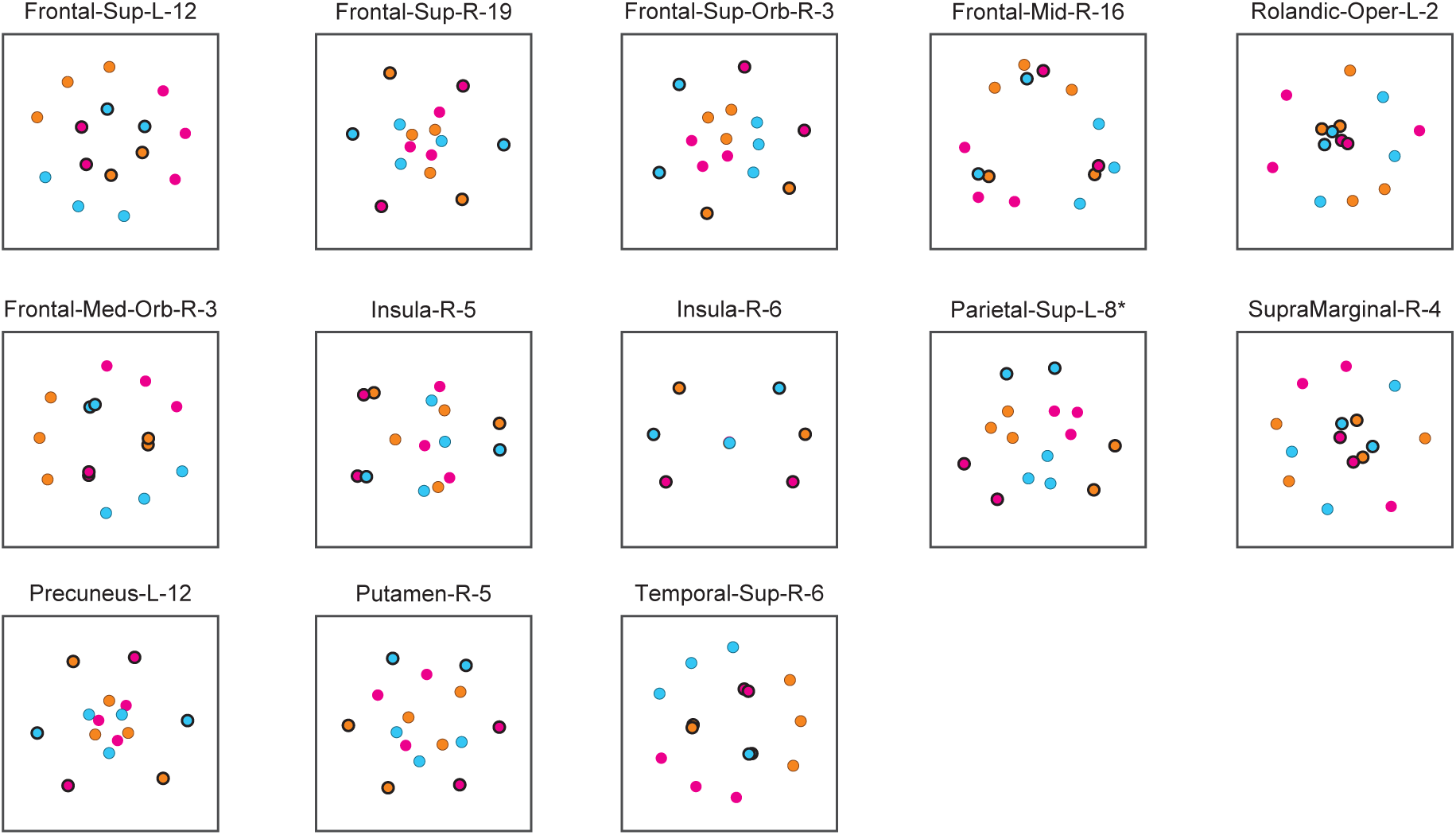
Representation of temporal community structure in negatively selective parcels. Multidimensional reduction of the pair-wise distance matrix averaged over path permutations and observers. Communities are distinguished by color and linking objects by a black outline. Thirteen parcels exhibited lower separability between communities than within communities (*t^BW^ <* 0). Identity-selective parcels are marked with *\**.

It is instructive to also compare parcels that were not classified as either identity- or community-selective. Results for fifteen randomly chosen ‘non-selective’ parcels are shown in **Supplementary Fig. S5**. Perhaps not surprisingly, the results were quite heterogeneous and few differences reached statistical significance when averaged over parcels (**Supplementary Fig. S6**). However, in several individual parcels, clustering by communities and/or prominent representation of linking objects was evident.

As a final control, we analyzed average pairwise distances in the responses obtained from unstructured sequences. Here we failed to observe significant deviations from the grand average distance, either for internal and linking objects or within and between communities (**Supplementary Fig. S6**). This corroborates that the results obtained from structured sequences were owed to the experience of temporal communities.

## 4 Discussion

We investigated incidental and automatic learning of regularities and dependencies without explicit behavioural task (Perruchet and Pacton, 2006; Aslin, 2017; Schapiro and Turk-Browne, 2015; Saffran and Kirkham, 2018; Perruchet, 2019; Fiser and Lengyel, 2022). Our aim was to compare the cortical basis of concurrent learning at two timescales, namely, learning of view-invariant object recognition at the timescale of object presentations (3 s) (Wallis and Bülthoff, 2001; Cox et al., 2005; Tian and Grill-Spector, 2015) and learning of task-irrelevant contingencies in the sequence of object presentations (“temporal communities” lasting ∼30 s) (Miyashita, 1988; Fiser and Aslin, 2002; Turk-Browne et al., 2005, 2009; Kakaei et al., 2021; Śaringer et al., 2022). Our results suggest that cortical representations of both object identity and of temporal community structure coexist in large parts of ventral occipitotemporal cortex, together with a prominent representation of the transition points between communities (linking objects).

Previous studies have localized view-invariant object representations in inferior temporal cortex (IT) and lateral occipital complex (LOC) (Śary et al., 1993; Grill-Spector et al., 2001; Van Meel and de Beeck, 2020). Single-unit responses in IT of non-human primates reflect the intrinsic contingencies of an invariant representation and correlate closely with recognition performance (Li and DiCarlo, 2008, 2010, 2012; Jia et al., 2021). Human fMRI show differential adaptation in IT for congruent and incongruent shapes (Van Meel and Op de Beeck, 2018). In addition, evidence for view-invariant representations has been reported in primary-visual cortex (Eger et al., 2008), at more anterior sites such as fusiform gyrus, and ventral occipito-temporal cortex (Brants et al., 2016; Visconti di Oleggio Castello et al., 2021), as well as in several areas of the dorsal pathway (Poirier et al., 2006; Konen and Kastner, 2008; Jeong and Xu, 2016; Freud et al., 2017; Visconti di Oleggio Castello et al., 2021).

Our results on cortical sites with view-invariant object representations confirm and extend these previous findings, as described in our companion study (Kakaei and Braun, 2023). In brief, multivariate representations of cross-validated object identity were established for smallish ‘functional parcels’ (∼ 1.7*cm*^3^ cortex volume) defined previously by a functional parcellation (MD758; Dornas and Braun 2018). Parcels in which significant identity information was prevalent (Allefeld et al., 2016) were located in both the ventral and dorsal visual pathways, beginning with early visual areas (V1-hV4), extending to more anterior parts of ventral occipitotemporal cortex into anterior inferior temporal cortex, as well as to anterior inferior frontal cortex. (Kakaei and Braun, 2023).

Our motivation to compare cortical representations of object shape and temporal object sequence stemmed from classical studies of object recognition in non-human primates (Miyashita, 1988; Erickson and Desimone, 1999). These studies showed that the responsiveness of single-neurons in inferotemporal cortex IT developed selectivitity not only for object identity, but additionally also for object sequence, after animals had repeatedly viewed the same visual objects in the same sequential order. The sequence representation exemplified incidental learning, because the sequential order was behaviourally irrelevant.

In extensive subsequent work with “paired-associate tasks”, the sequential order of objects was made task-relevant so that learning of temporal associations became explicit. Over the course of training, the prevalence of pair-encoding neurons was found to increase in anterior parts of inferiotemporal cortex IT Messinger et al. 2001; Naya et al. 2001, 2003; Hirabayashi and Miyashita 2014. Additionally, neurons in IT were found to encode “object-general semantic value” in the sense of identifying whether a particular object is “familiar” or “novel” (Tamura et al., 2017). We wondered whether such “object-general” information could extend to membership in a “temporal community” of objects.

Previous behavioural studies have shown that humans can implicitly learn spatiotemporal associations between objects and use these regularities to enhance their cognitive performance. Observers can automatically capture spatial (Fiser and Aslin, 2001) and temporal (Fiser and Aslin, 2002; Turk-Browne et al., 2008) regularities as both joint and conditional probabilities of stimuli co-occurrence. This surpasses simple object-object associations and extends to higher-order association probabilities, over multiple objects. Even when the conditional probability between object pairs are uniform and thus uninformative of the underlying association between objects, humans are sensitive to higher-order regularities (higher-moments of conditional probability distribution) (Schapiro et al., 2013; Karuza et al., 2017b; Kahn et al., 2018; Kakaei et al., 2021). This capability for incidental learning of complex regularities can facilitate performance in various domains, including language(e.g. Saffran et al. 1996), motor(e.g. Hunt and Aslin 2001), spatial attention(e.g. Chun and Jiang 1998; Jiang and Wagner 2004), and object recognition learning(e.g. Kakaei et al. 2021).

The literature on implicit or explicit learning of temporal associations shows that both domain-specific and domain-general brain regions can be involved (for reviews see Batterink et al. 2019; Fiser and Lengyel 2022). Neural correlates of statistical learning are evident in early domain-specific sensory areas where spatio-temporal regularities are first extracted, to mid-level sensory areas where these representations are supposedly integrated. In the visual domain, spatio-temporal regularities emerge in lateral and ventral occipito-temporal and parieto-occipital regions in humans (Turk-Browne et al., 2009; Rosenthal et al., 2016; Karuza et al., 2017a; Henin et al., 2021) and is observed in inferiotemporal and anterior inferiotemporal regions in non-human primates (Miyashita, 1988; Sakai and Miyashita, 1991; Meyer et al., 2014; Kaposvari et al., 2018). More abstract and generalized representation of temporal associations have been reported in more down-stream, domain-general areas such as MTL, striatum and frontal regions. Moreover, the majority of studies points to an to essential role of medial temporal lobe (MTL), particularly hippocampus, in statistical learning (Schendan et al., 2003; Turk-Browne et al., 2009, 2010; Schapiro et al., 2012, 2013, 2016; Schapiro and Turk-Browne, 2015; Hsieh et al., 2014; Hindy et al., 2016). This is particularly true when sequences are repeated and an ordinal knowledge is of particular interest (for reviews see Davachi and DuBrow 2015; Eichenbaum et al. 2016). MTL seems to be engaged in statistical learning occur early in the learning process, but seems to disengage as learning progresses, particularly after consolidation. Concurrently, the encoding of statistical knowledge seems transfer from MTL to the striato-frontal network (Durrant et al., 2013; Batterink et al., 2019). Higher cortical regions in insular cortex and prefrontal cortex (PFC), including inferior frontal gyrus (IFG) and medial prefrontal cortex (mPFC), also show sensitivity to statistical regularities, particularly when the complexity increases (Schapiro et al., 2013; Wang et al., 2017; Giorgio et al., 2018; Karlaftis et al., 2019; Kourtzi and Welchman, 2019; Henin et al., 2021).

Here we followed Schapiro and colleagues (Schapiro et al., 2013) and used object sequences with higher-order “temporal community structure”. In such sequences, pair-probabilities are uniform in that every object is succeeded by one of four other objects with equal probability. This avoids the novelty/surprise effects that would arise if some objects transitions had been more common/rare than others. We term sequences with temporal communities “strongly-structured”, in order to distinguish them from “unstructured” pseudo-random sequences where every object can be succeeded by any other object (Kakaei et al., 2021).

We studied cortical representations with a “representational similarity analysis” (RSA) approach. This approach relies on comparing pairwise distances of multivariate BOLD signal patters to objects, for example within and between communities. However, multivariate BOLD patterns are known to be significantly autocorrelated over 10s of seconds (Henriksson et al., 2015; Alink et al., 2015), in part due to haemodynamic effects (Friston et al., 1994; Zarahn et al., 1997). Accordingly, it is essential to distinguish between response similarity owing to “temporal community” effects and similarity that is merely due to temporal proximity (short latency) (Gilron et al., 2016; Cai et al., 2019). We took two measures to control for this confound and to distinguish between community and latency effects. First, we computed and analysed ‘residual distances’ by subtracting from each observed distance at a certain latency the *average* distance at that latency (see Methods 2.3.2). Second, we assessed consistency by analysing and comparing distances in different latency ranges, for example including or excluding short latencies. These measures turned out to be essential, as nearly the entire brain would have spuriously appeared to be ‘community-selective’ without them. The same measures also proved to be effective, as they revealed ‘community-selectivity’ only in multivariate BOLD responses to “strongly-structured” sequences and not in responses to “unstructured” sequences. Accordingly, we are confident that these measures identify genuine cortical representations of “temporal community”.

Our analysis of multivariate BOLD responses in 758 ‘functional parcels’ revealed two distinct kinds of ‘community-selectivity’. The first kind – termed *positively-selective* – showed *greater* similarity of responses within communities then between communities and was observed mostly in domain-specific brain regions. The second kind – termed *negatively-selective* – exhibited *lesser* similarity of responses within communities and was observed mostly in domain-general areas. Below we discuss these functionally and anatomically distinct groups of parcels in more detail.

Positively community-selective parcels showed a domain-specific alternation of neural representation with “temporal community” in early- and mid-level visual areas. Enhanced similarity within temporal communities was particularly evident in the ventral pathway, specifically in visual areas V1-hV4, inferior and lateral occipito-temporal networks. This representation of higher-order statistical regularities was a subset of the cortical representation of object identity. While it was much narrower (comprised fewer parcels), it covered nearly the same extent of the ventral visual pathway. This is consistent with earlier findings that early and mid-level visual areas are sensitive to temporal regularities and can flexibly alter their activity pattern to represent the temporal context. It is also consistent with the classical observation that representations of temporal association develop conjointly with representations of object identity (Miyashita, 1988; Erickson and Desimone, 1999).

In contrast to earlier studies cited above, we failed to observe positive community-selectivity in the medial temporal lobe (MTL). MTL engages early in the learning process and after consolidation, the engram gets transferred to the striatum. As our observations cover multiple days, overnight consoliation could have occurred already after the first session, which might explain the absence of community representation in MTL (conversely the presence in putamen). As our analysis was ‘data-hungry’, observations from all sessions had to be combined. Accordingly, the present study might simply have lacked the statistical power to identify community-selectivity in MTL during the early stages of learning.

Taken together, these results suggest that incidental learning of temporal associations at both shorter and longer time-scales occur at all levels of the ventral visual pathway. Both learning of view-invariant representations of visual objects (on a time-scale of seconds) and learning of temporal contingencies in the object sequence (on a time-scale of tens of seconds) are represented at overlapping locations along this pathway. This functional overlap implies the the visual hierarchy employs *convergent representations* (Grill-Spector and Weiner, 2014) that integrate information from a range of time-scales and that can potentially mediate context-dependent enhancement of recognition performance (Kakaei et al., 2021).

We also identified negatively community-selective parcels in domain-general functional networks, including superior parieto-frontal, ventral attention, and fronto-striatal networks, that have been reported to be engaged by different implicit learning paradigms (Batterink et al., 2019). As these parcels were not only negatively community-selective, but additionally lacked significant representation of object identity, their community representation would seem to serve an altogether different purpose than the representation in visual areas, which was mostly positive and identity-selective.

Previous studies provide ample evidence for more abstract functions of the implicated locations in domain-general cortex. Areas such as prefrontal cortex (PFC) are thought to reflect higher-order statistics of event (Henin et al., 2021) and decision strategies adopted by observers (Wang et al., 2017; Giorgio et al., 2018; Karlaftis et al., 2019; Kourtzi and Welchman, 2019). Insula and inferior frontal are thought to be engaged by working memory tasks, especially under conditions of high load (Rottschy et al., 2012) and to contribute to goal-directed behaviour by interacting with the medial temporal lobe hippocampus (Rusu and Pennartz, 2020). Orbitofrontal cortex (OFC) is thought to be engaged when more abstract representations or ‘cognitive maps’ are required (Wilson et al., 2014; Schuck et al., 2016; Christophel et al., 2017; Behrens et al., 2018; Rusu and Pennartz, 2020; Knudsen and Wallis, 2022).

Cognitive maps could be context-specific and individuate objects in the current community, without necessarily identifying either the community or objects in other communities. When the context changes (because the sequence visits another community) the same map could transform to individuate objects in the new community. This would be analogous to invariant response patterns of grid-cells in different environments observed in the entorhinal cortex of rodents and humans (Fyhn et al., 2007; Doeller et al., 2010; Constantinescu et al., 2016). Such a context-specific map would reconcile the relative dissimilarity of response patterns within communities and the relative similarity between communities, which is characteristic of negatively community-selective parcels (**Fig. S6**) in our study and which previous studies have observed in medial prefrontal cortex (mPFC) (Schapiro et al., 2013).

An alternative account of negative community-selectivity could invoke the comparative salience of linking objects and the transition to a new community. While the object sequence remains within a community, observers can expect to see further objects from this community. Transitions to a new community violate this expectation which could be experienced as a salient event and could generate a ‘novelty’ (for a review see Uddin 2015). In fact, our results confirm the relative dissimilarity of linking object to other objects of the same community (and relative similarity to other objects in different communities) (**Fig. S6**). However, as we took care to establish that these effects that hold over all latencies (not only over short latencies), this alternative account cannot fully explain our findings.

Finally, we note that the behavioural task in our state created no behavioural benefit for maintaining individual object representations. Observers were required merely to categorize objects as ‘familiar’ or ‘not familiar’, and were not required to identify individual objects ‘by name’. This could have favoured the absence of object-identity representations in domain-general areas, irrespective of the exact mechanism behind negative community-selectivity.

## Data and code availability

Direct linear discriminant analysis and prevalence inference is available on github.com/cognitive-biology/DLDA. MR data will be made available upon request.

## Credit authorship contribution statement

**E. Kakaei:** Conceptualization, data curation, formal analysis, visualization, writing of original draft. **J. Braun:** Conceptualization, linear algebra, formal analysis, supervision, reviewing & editing.

## Acknowledgments

We thank Claus Tempelmann, Martin Kanowski and Denise Scheermann at the Magnetic Resonance Imaging Laboratory of the Department of Neurology of Otto-von-Guericke University, Magdeburg. We also thank Stepan Aleshin for helpful discussions and constructive comments. This study was funded by the federal state Saxony-Anhalt and the European Structural and Investment Funds (ESF, 2014-2020), project number ZS/2016/08/80645, as part of doctoral program ABINEP (Analysis, Imaging and Modelling of Neuronal Processes).

## A Appendices

**Table A.1:** List of community selective parcels and their anatomical region. Numerical parcel ID, geometrical centroid x/y/z in MNI and between-community separability *t^BW^*.

| Region | Parcel No. | MNI | | | $t^{BW}$ |
| --- | --- | --- | --- | --- | --- |
|  |  | x | y | z |  |
| Middle frontal | 109 | 48 | -3 | 56 | 2.9 |
| Calcarine | 337 | -12 | -99 | -5 | 5.2 |
|  | 344 | 9 | -85 | 6 | 5.2 |
| Lingual | 363 | -12 | -65 | -5 | 3.8 |
| Occipital (middle) | 400 | -38 | -86 | 4 | 3 |
|  | 405 | -16 | -100 | 1 | 6.3 |
| Occipital (inferior) | 424 | -31 | -83 | -8 | 4.5 |
|  | 428 | 36 | -85 | -7 | 5.9 |
|  | 430 | 42 | -73 | -9 | 3.6 |
| Fusiform | 432 | -27 | -71 | -11 | 4.9 |
|  | 436 | -31 | -53 | -13 | 4.1 |
| Postcentral | 483 | 51 | -26 | 47 | 3.8 |
| Temporal (inferior) | 751 | 51 | -67 | -8 | 5.1 |
|  | 755 | 46 | -53 | -11 | 5.5 |
| Superior frontal | 45 | -23 | 58 | 23 | -4.1 |
|  | 69 | 21 | 63 | 4 | -6.4 |
| Superior frontal (orbital) | 77 | 23 | 61 | -5 | -6.8 |
| Middle frontal | 118 | 44 | 35 | 34 | -3.8 |
| Rolandic operculum | 187 | -42 | -25 | 18 | -3.2 |
| Superior frontal (medial orbital) | 248 | 9 | 65 | -9 | -4.6 |
| Insula | 270 | 39 | 20 | -4 | -3.6 |
|  | 271 | 41 | 7 | 4 | -3.8 |
| Parietal (superior) | 499 | -28 | -69 | 50 | -4.2 |
| Supramarginal | 540 | 56 | -44 | 29 | -4.4 |
| Precuneus | 570 | -3 | -62 | 46 | -3.5 |
| Putamen | 619 | 26 | 11 | 6 | -4.2 |
| Temporal (superior) | 650 | 63 | -8 | 5 | -3.5 |

## Supplement

**Figure S1:**
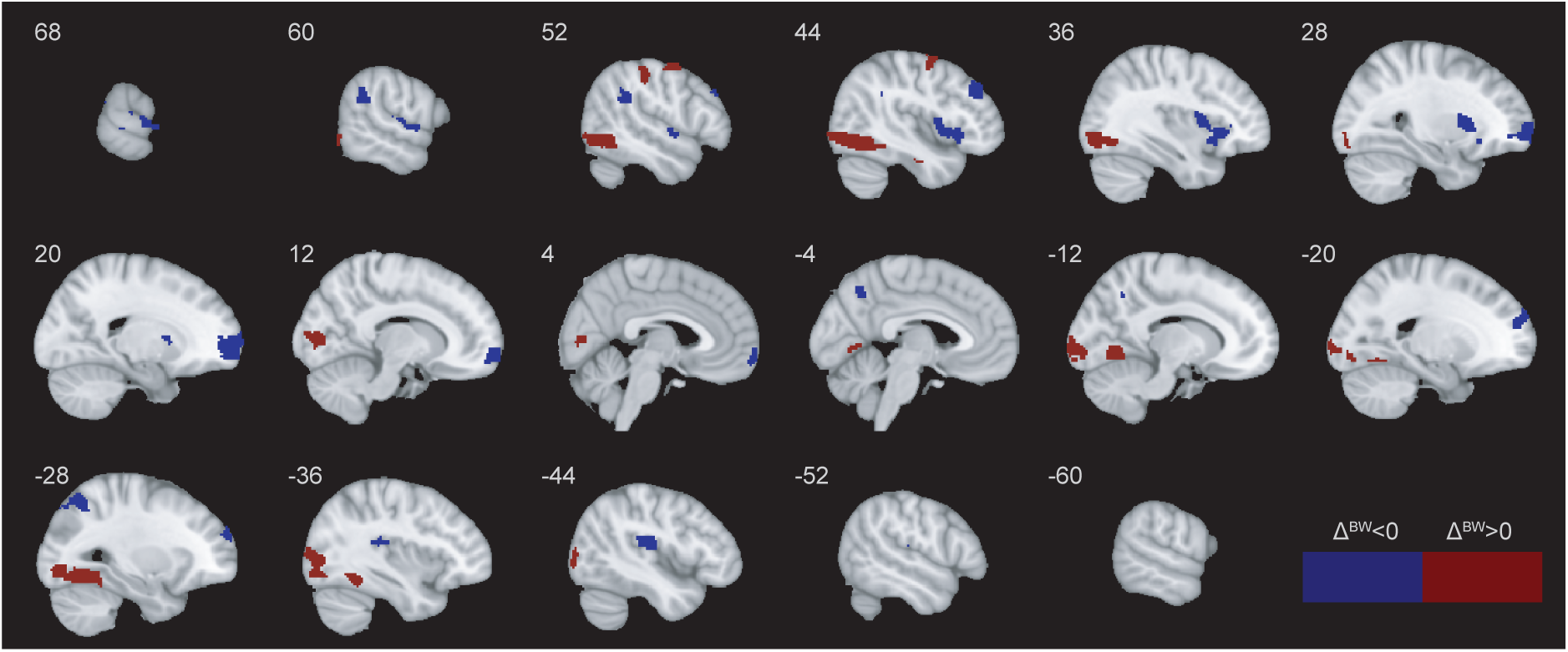
Neural representations of temporal community structure. Community-selective parcels in red have significantly higher separability between communities than within *t^BW^ >* 0, and the other way around for blue parcels *t^BW^ <* 0. Sagittal slices (8 mm thickness) range from *X* = −60 to *X* = +68 (MNI).

**Figure S2:**
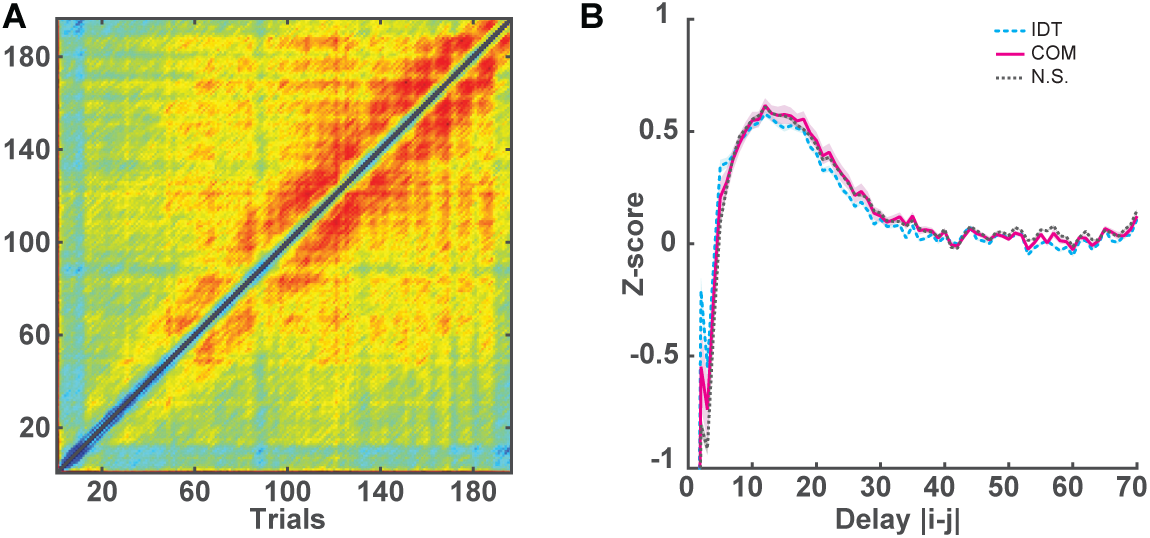
Delay-dependence of the distance between response pairs. Pairwise euclidean distance *dw,u,r* (*i, j*) between responses *i* and *j* was computed for each parcel *w*, run *r*, and subject *u*. For each parcel *w*, the distance matrix *Tw* (*i, j*) = ⟨*dw,u,r* (Δ*i*)⟩*u,r* was averaged over observers and runs. **A)** Distance matrix *T* (*i, j*) = ⟨*Tw* (*i, j*)⟩*w* averaged over parcels. Note that delay-dependence grows over the duration of a run. **B)** Average delay-dependence *T* (Δ*i*) = ⟨*dw,u,r* (Δ*i*)⟩*w,u,r* – in terms of z-score ± S.E.M over parcels – for three groups of parcels: identity-selective (cyan), community selective (magenta) and non-selective parcels (black). Adapted from Kakaei and Braun 2023.

**Figure S3:**
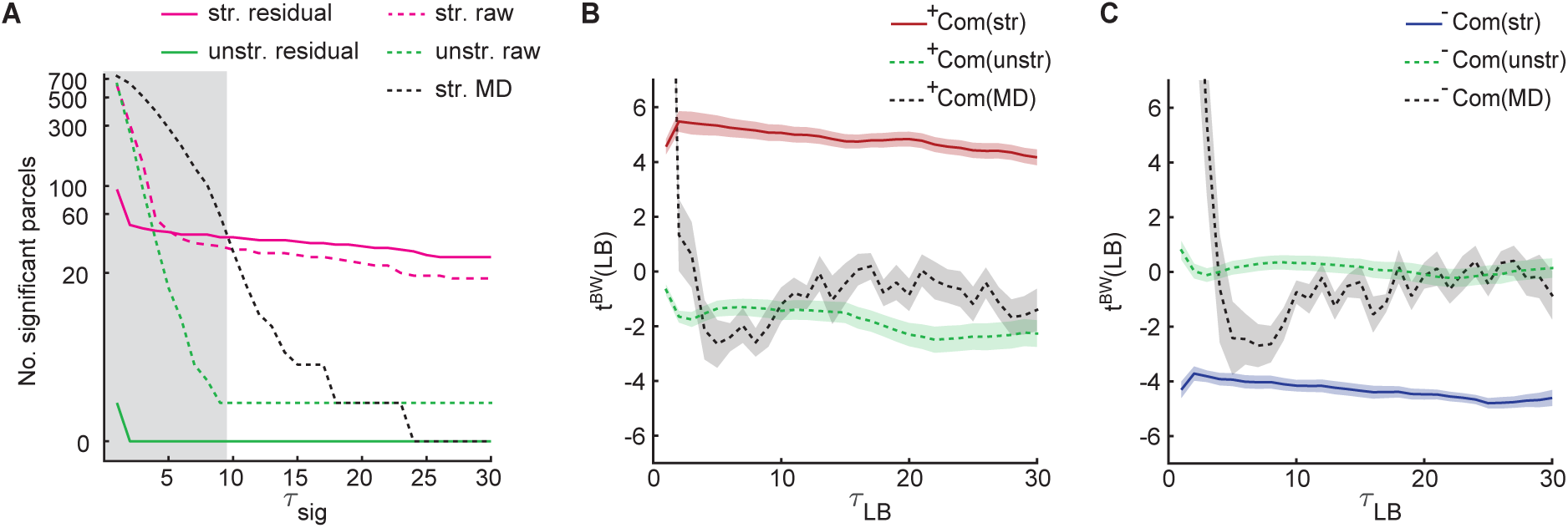
Comparison of controls for community representation. **A)** Apparent community selectivity, as measured by separability *t^BW^* that is consistently significant for all latency ranges with lower bounds *τ_LB_* ∈ {1, 2*, . . ., τ_sig_* }. The number apparently selective of parcels is shown for latency-corrected, residual distances (solid) and for non-latency corrected, raw distances (dashed). Distances were obtained from structured sequences (magenta), from unstructured sequences(green), as well as from static, average distance matrices (black). **B)**Between community separability *t^BW^* ± *S.E.M.*, as function latency lower bound *τ ^BW^*), for *positively* community selective parcels, based on structured sequences (solid red), unstructured sequences (dashed green), and static average distance matrices (dashed black). **C)** Between community separability *t^BW^* calculated for *negatively* community selective parcels, based on structured sequences (solid blue), unstructured sequences (dashed green), and average distance matrices (dashed black).

**Figure S4:**
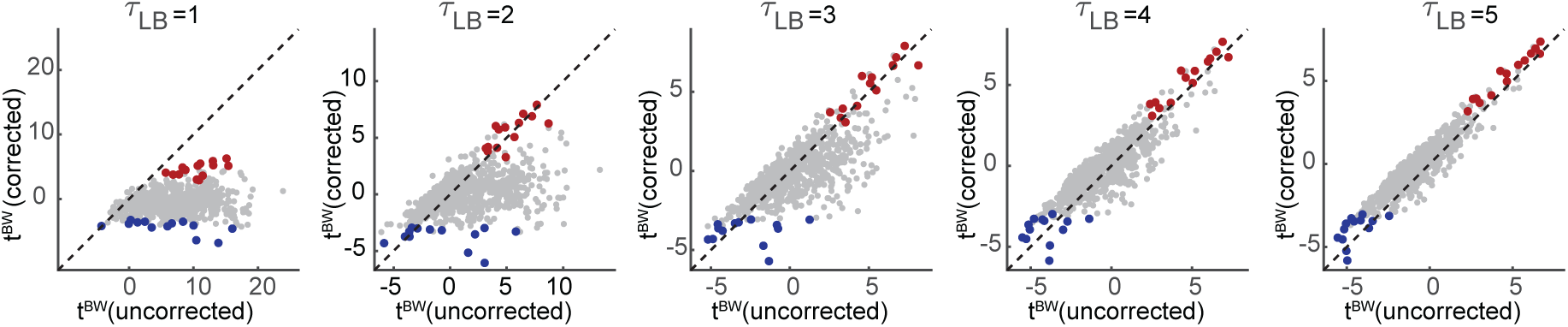
Community selectivity, latency bound and correction for temporal correlations. Comparison of between-community separability *t^BW^* for uncorrected and for corrected distances, as well as for different latency lower bounds *τ_LB_* . Parcels classified as *positively* (Δ*^BW^ >* 0) or *negatively* (Δ*^BW^ <* 0) community-selective are shown in red and blue, respectively. parcels with higher separability for between-community pairs (Δ*^BW^>* 0) and for within-community pairs (Δ*^BW^<* 0) are shown in red and blue, respectively. Dashed lines indicate identity.

**Figure S5:**
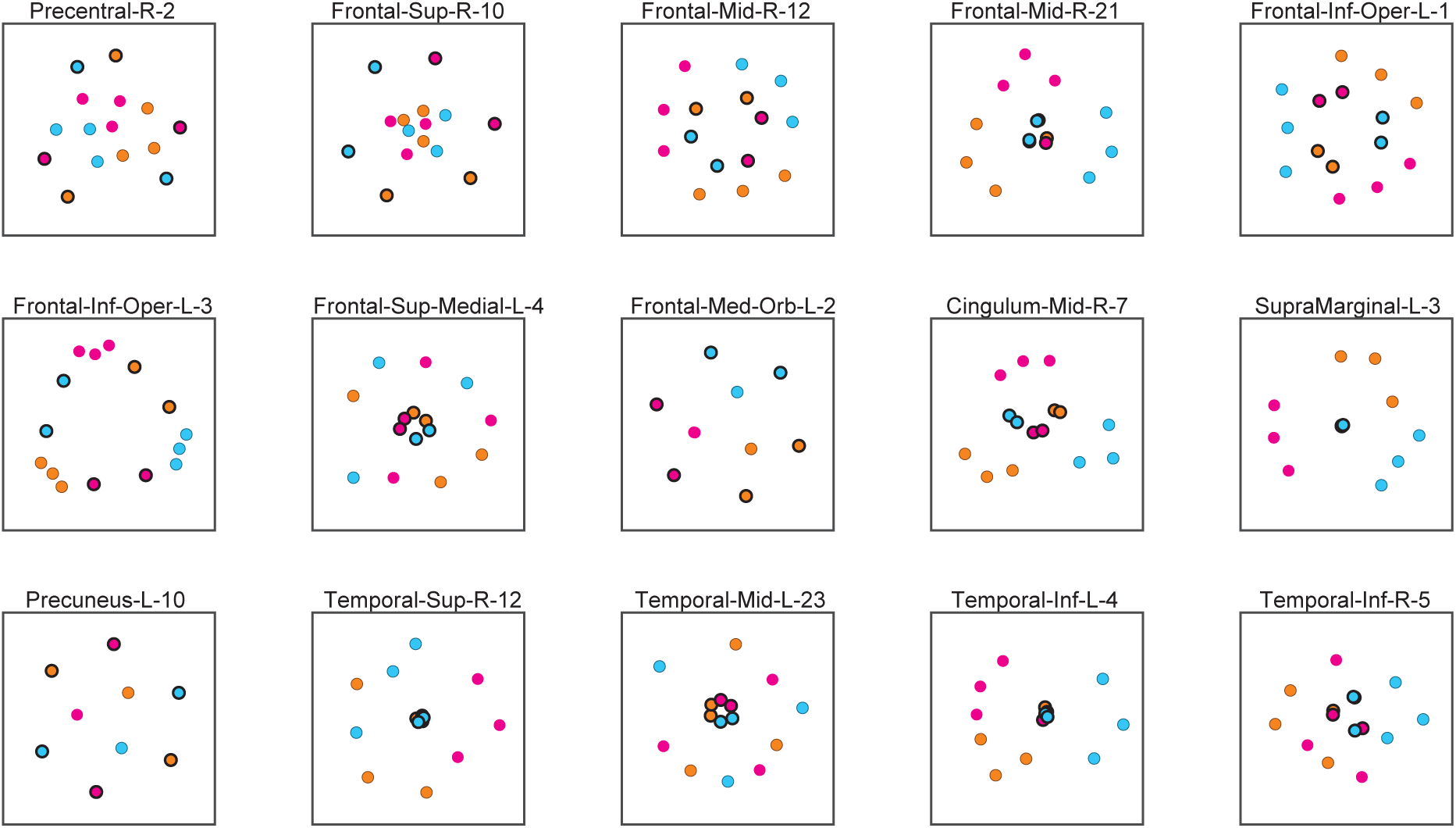
Representation of temporal community structure in other, non-selective parcels. Multidimensional reduction of the pair-wise distance matrix, averaged over path permutations and observers. Communities are distinguished by color and linking objects by a black outline. The fifteen parcels shown were selected randomly from non-indentity-and non-community-selective parcels (*t^BW^* ≈ 0).

**Figure S6:**
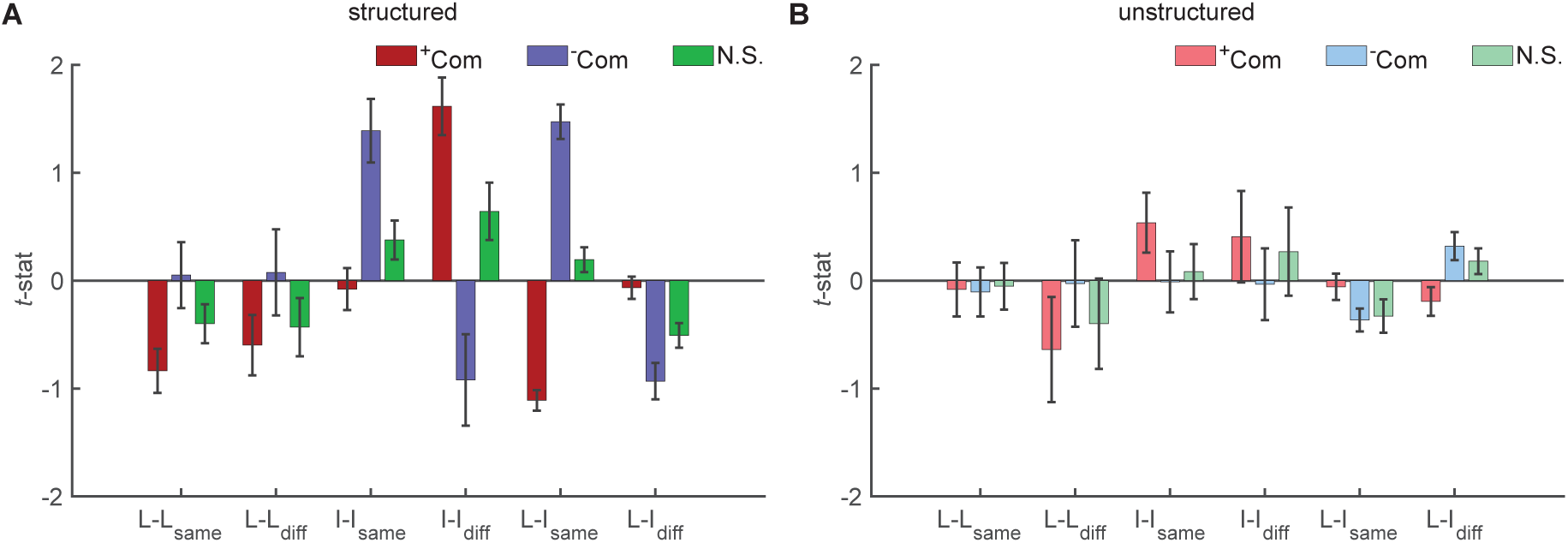
Separability of different types of object pairs, composed of linking (L) and/or internal (I) objects, within the same or different communities, for three different sets of parcels: *positively* community selective parcels (red) (compare Fig. 6), *negatively* community selective parcels (blue, compare Fig. 7), and randomly chosen, non-selective parcels (green, compare **Fig. S5**). **A)** Average separability *t* ± *S.E.M* for each group of parcels computed from responses to structured sequences. **B)** Average separability *t*±*S.E.M* for each group of parcels computed from responses to unstructured sequences.

